# CHARMM-GUI Multicomponent Assembler for Modeling and Simulation of Complex Multicomponent Systems

**DOI:** 10.1101/2023.08.30.555590

**Authors:** Nathan R. Kern, Jumin Lee, Yeol Kyo Choi, Wonpil Im

**Affiliations:** Department of Computer Science & Engineering, Lehigh University, Bethlehem, PA, USA; Department of Biological Sciences, Lehigh University, Bethlehem, PA, USA; Departments of Biological Sciences, Chemistry, Bioengineering, and Computer Science & Engineering, Lehigh University, Bethlehem, PA, USA

## Abstract

Atomic-scale molecular modeling and simulation are powerful tools for computational biology. However, constructing models with large, densely packed molecules, non-water solvents, or with combinations of multiple biomembranes, polymers, and nanomaterials remains challenging and requires significant time and expertise. Furthermore, existing tools do not support such assemblies under the periodic boundary conditions (PBC) necessary for molecular simulation. Here, we describe *Multicomponent Assembler* in CHARMM-GUI that automates complex molecular assembly and simulation input preparation under the PBC. We demonstrate its versatility by preparing 6 challenging systems with varying density of large components: (1) solvated proteins, (2) solvated proteins with a pre-equilibrated membrane, (3) solvated proteins with a sheet-like nanomaterial, (4) solvated proteins with a sheet-like polymer, (5) a mixed membrane-nanomaterial system, and (6) a sheet-like polymer with gaseous solvent. *Multicomponent Assembler* is expected to be a unique cyberinfrastructure to facilitate innovative studies of complex interactions between small (organic and inorganic) molecules, biomacromolecules, polymers, and nanomaterials.

## Introduction

Molecular dynamics (MD) simulation becomes essential to study diverse molecular phenomena at atomic resolutions. Advances in computational power and algorithms have enabled simulation models with atom counts in the range of 100 million to several billion, including studies of metal nucleation and sliding^1–4^, bacterial cytoplasm^5–7^, eukaryotic gene loci^8^, synaptic bouton^9^, and viral capsids^10,11^. Preparing MD simulation systems typically requires determining the model size and composition, solving an NP-hard packing problem, and having topological information of each molecule and associated force field (FF) together with simulation input parameters. However, each of these steps presents a challenging barrier for researchers attempting to enter the field, including many experimental researchers who seek to supplement their studies with molecular modeling and simulation.

Many software applications have been developed to facilitate various steps of atomistic model preparation, including FFParam^12^, FFTK^13^, SwissParam^14^, Antechamber^15^, CGenFF^16–18^, MATCH^19^, OpenFF^20^, and CHARMM-GUI *Ligand Reader & Modeler*^21^ for FF preparation; PACKMOL^22–24^, cellPACK^10,11^, LipidWrapper^25^, and Soup^26^ for molecular packing; and *FF-Converter*^27,28^ and ParmEd^29^ for preparing and converting inputs for several simulation programs. However, except for a few notable model types, no free software package can handle all steps of model preparation without significant experimental knowledge, manual intervention, or use of *ad hoc* scripts. Certain common molecular configurations are already handled by simulation and analysis programs directly, such as AmberTools^30^, GROMACS^31,32^, NAMD/VMD^33,34^, and OpenMM^35^, which can add a box of water and ions to a single molecule of interest or to a pre-arranged molecular complex or mixture. Similarly, several tools exist to facilitate building combined protein-membrane models by using the template copying method for water placement and by using either a membrane template scheme for positioning lipids or solving the packing problem for the membrane-protein only^36–42^. In each of these cases, only a single protein or pre-oriented protein complex can be handled per molecular model. Furthermore, while PACKMOL, cellPACK, and LipidWrapper can pack a wide variety of molecules into nearly any defined shape, neither program currently handles the periodic boundary conditions (PBC) necessary for MD simulations. In addition, significant post-processing is typically required to use the output of these tools to prepare inputs for different simulation programs.

CHARMM-GUI (https://www.charmm-gui.org)^43^ is a cyberinfrastructure that guides researchers through building simulation-ready atomistic and coarse-grained models containing a single component or complex of interest in solution or membrane environments. Recent developments have also extended its functionality to cover various nanomaterials and polymer models^44,45^. In this work, we present CHARMM-GUI *Multicomponent Assembler* (MCA) that solves a complex packing problem for many components of interest under the PBC, enables using pre-equilibrated membrane or membrane-like (nanomaterials and polymer) materials, and generalizes the template-based solvent building approach to cover non-water and mixed solvents. To illustrate MCA’s versatility for heterogeneous system building, we prepared and simulated 6 challenging systems at various densities, for a total of 20 systems, using only components generated by other CHARMM-GUI modules: (1) solvated proteins, (2) solvated proteins with a pre-equilibrated membrane, (3) solvated proteins with a sheet-like nanomaterial, (4) solvated proteins with a sheet-like polymer, (5) a mixed membrane-nanomaterial system, and (6) a sheet-like polymer with gaseous solvent.

## Results and Discussion

### 1. Workflow of *Multicomponent Assembler*

MCA handles molecular components by grouping them into 5 categories (**Table 1**) that differ by their general positioning requirements and assumptions. MCA combines components into 6 overall molecular configurations (**Figure 1**) using the workflow described below and in **Figure 2**. Note that MCA currently uses the CHARMM36(m) FF^46^ for protein and lipid components, CGenFF^16– 18,47,48^ for polymers, and INTERFACE FF^49–54^ for nanomaterials.

**Table 1.**
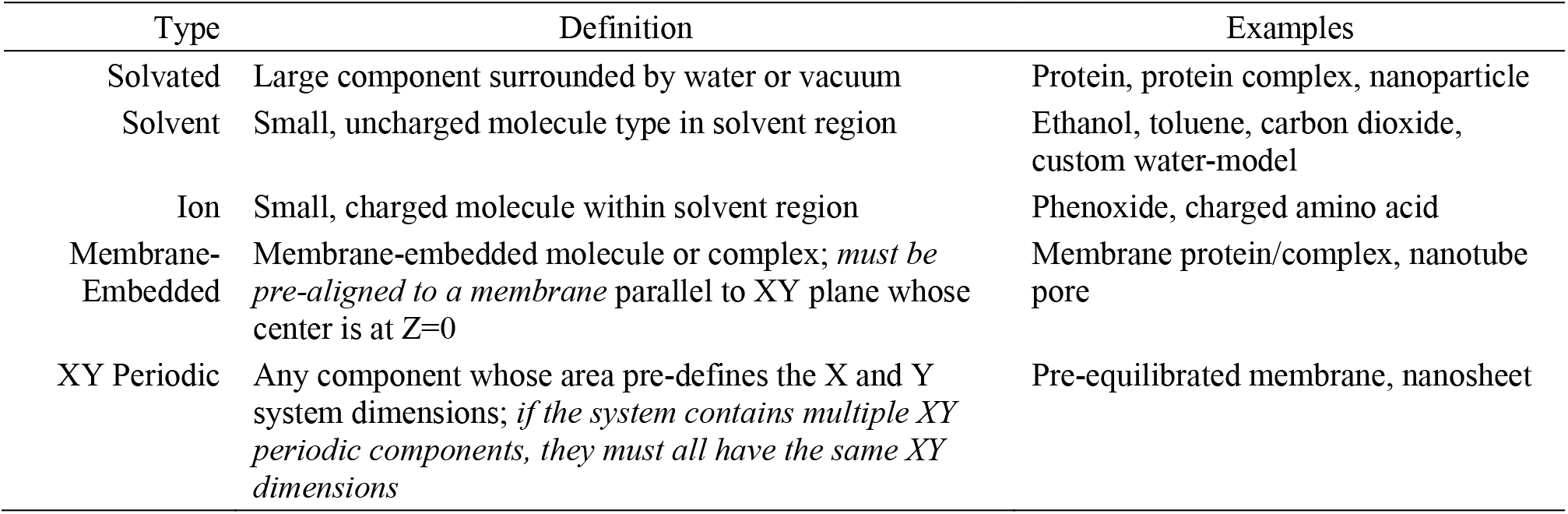
Supported component types. Each uploaded component must have a component type based on its positioning and packing requirements.

**Figure 1.**
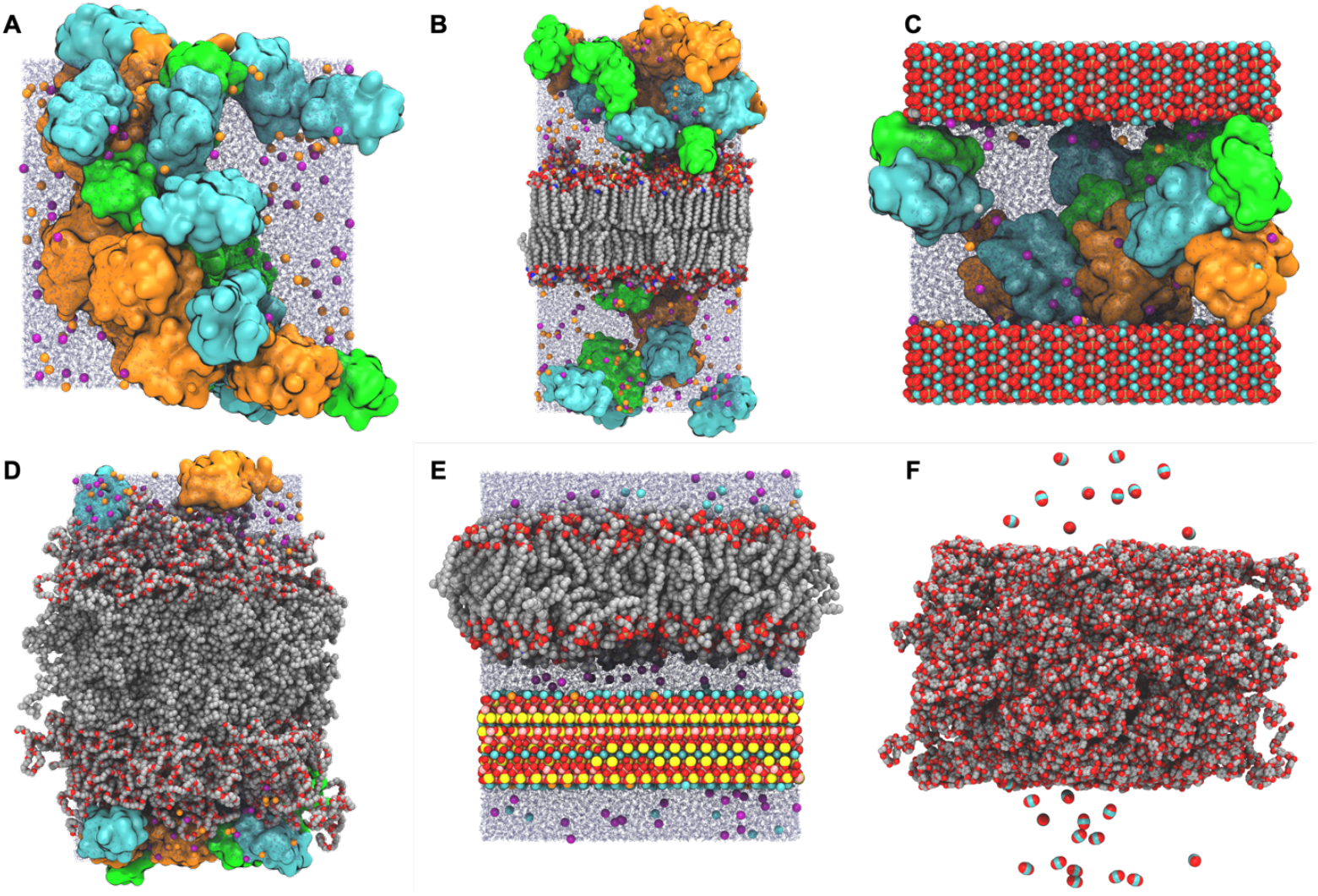
Example system configurations generated by *Multicomponent Assembler*. In all figures containing protein, ubiquitin (PDB: 1UBQ) is orange, villin (PDB: 1VII) is green, and protein G (PDB: 3GB1) is cyan. K^+^ and Cl^-^ ions are shown as orange and purple spheres, respectively, and water molecules as blue translucent lines. When included, KCl concentration is 0.15 M. (A) Proteins with water and ions only. (B) Pre-equilibrated axolemma membrane combined with proteins. (C) Proteins with a hydroxyapatite slab centered on the unit cell’s Z boundary. (D) Artificial membrane of polyethylene oxide-poly(ethylethylene) (EO_40_EE_37_) with proteins. (E) Supported lipid bilayer consisting of POPC membrane and mica separated by a 20 Å water layer. (F) CO_2_ adsorption on a polyethylene terephthalate (PET) membrane.

**Figure 2.**
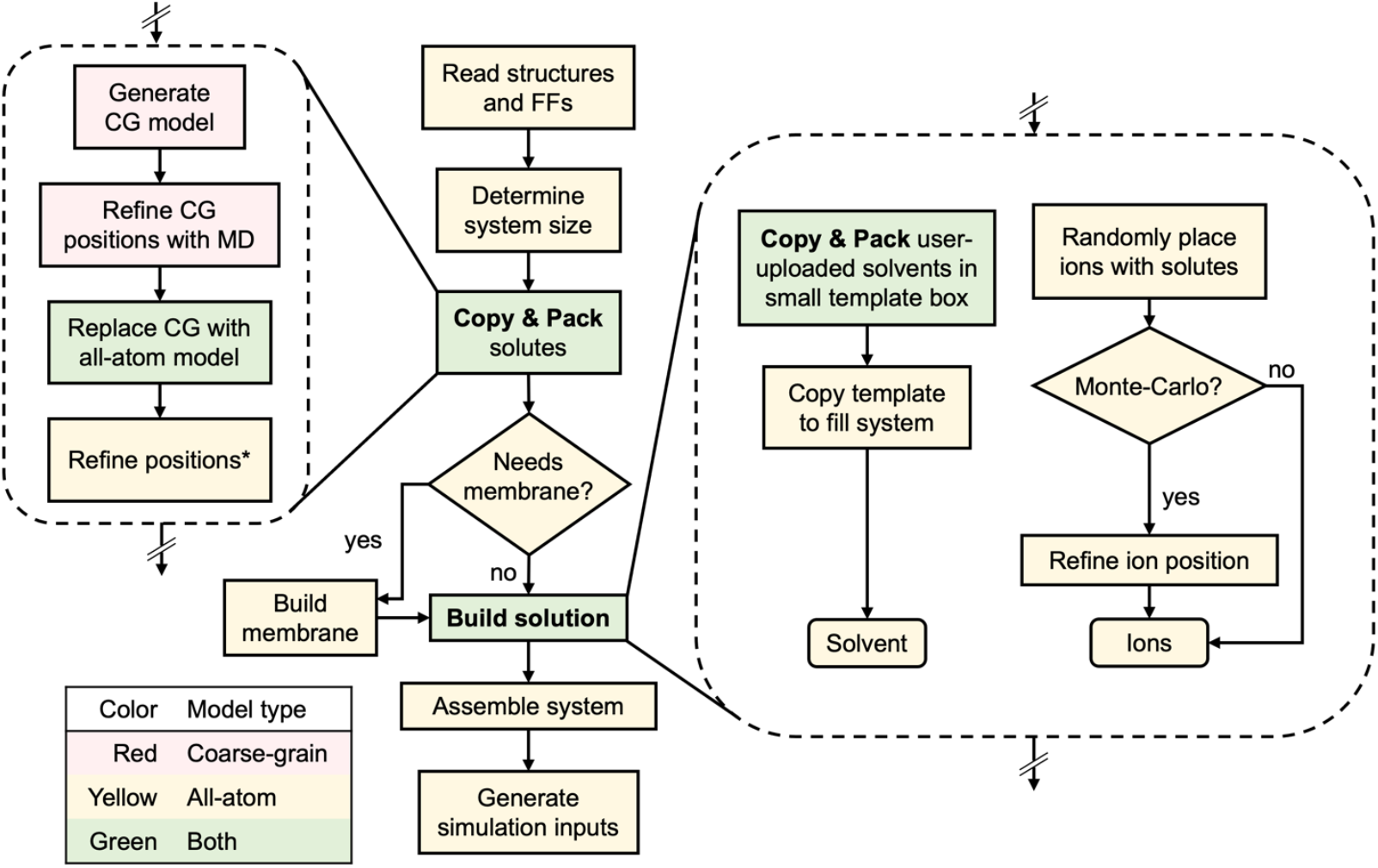
*Multicomponent Assembler* workflow. Step colors indicate which model types they use. The Copy & Pack procedure is used to place solutes and to create a small solvent template. ^*^Positions are refined using the algorithm in Supporting Information **Algorithm 1**.

#### STEP 1 – Read structures and FFs

Because CHARMM-GUI uses CHARMM for model building and manipulation, it is required to provide molecular structures in both CHARMM protein structure file (PSF) and coordinate (CRD) formats. Additionally, for periodic (nanomaterial) structures with image bonds (between the primary system and the PBC systems), listing the bonds in a CHARMM-formatted image PSF file is necessary. Molecules containing residues already present in the CHARMM and INTERFACE FFs are recognized automatically, but any additional residue topologies and FFs must be provided in CHARMM’s residue topology file (RTF) and parameter (PRM) formats with unique identifiers that do not conflict with the existing FFs. Note that these files can be obtained from other CHARMM-GUI modules such as *PDB Reader & Modeler*^55^, *Glycan Reader & Modeler*^56^, *Ligand Reader & Modeler*^21^, *Nanomaterial Modeler*^44^, and *Polymer Builder*^45^.

For each uploaded component, MCA uses CHARMM internal functions to determine the molecular dimensions, volume, solvent-accessible volume, mass, charge, number of residues, and radius of gyration. To facilitate membrane building, the uploaded coordinates are used to determine the volume of the molecular regions that would be located within, above, and below a membrane centered at Z=0 with a hydrophobic thickness of 24 Å. The component’s dimensions are calculated as its bounding box when the component’s primary, secondary, and tertiary axes are aligned with the X, Y, and Z axes, respectively. The component’s length is determined as the maximum distance between any two atoms within the structure. This value is needed to estimate the minimum box size that prevents any component from interacting with its own PBC images. The molecular volume is calculated by polling coordinates using a grid spacing of 0.5 Å within any atom’s van der Waals radius, with a maximum probe range of 6 Å and counting the empty holes within a molecule toward its volume. For solvent-accessible volume, the procedure is repeated with atomic radii increased by 1.4 Å, corresponding to the water radius.

To ensure that segment identifiers are both unique and meaningful, all segments from each input structure are written to separate PDB files to determine the segment type (protein, DNA, RNA, heterogen, carbohydrate, or water). Each input segment is then re-written to a PSF such that the first letter corresponds to the segment type (P, D, R, H, C, W), and the next two letters designate a unique ID. Thus, the first input protein segment is renamed to PAA, the 27^th^ to PBA, etc. A human-readable map of input-to-output segment IDs is written to a file (rename_map.txt) for the user’s convenience.

#### STEP 2 – Determine system size and positional constraints, and pack solutes

To properly specify the components in a given system, users must identify the type of each component as listed in **Table 1**. If there are no membrane-embedded components, users must specify whether a new membrane should be generated.

In the simplest case, the user already knows the exact number of copies of each component and system dimensions. However, commonly only the relative ratio of components with respect to each other is known. Similarly, instead of knowing the exact system dimensions, the user may know only the fraction of available volume that should be occupied by uploaded components (not including solvent and ions that are handled later). To guide in determining an appropriate system size and number of components, one can specify the relative component ratios, and specify either exact system dimensions or volume fraction and an approximation of the other quantity, as shown in **Figure 3**. MCA recommends the closest match to the approximated quantity with the following constraints: (1) no dimension is small enough that a component can interact with its own PBC images, and (2) the volume fraction is low enough that packing can plausibly succeed. To quickly estimate an upper bound on the maximum packing density, we rely on a heuristic that empty space should be no less than the total solvent accessible volume of all components. However, in our observation, achieving volume fractions higher than 30% usually requires significant trial-and-error.

**Figure 3.**
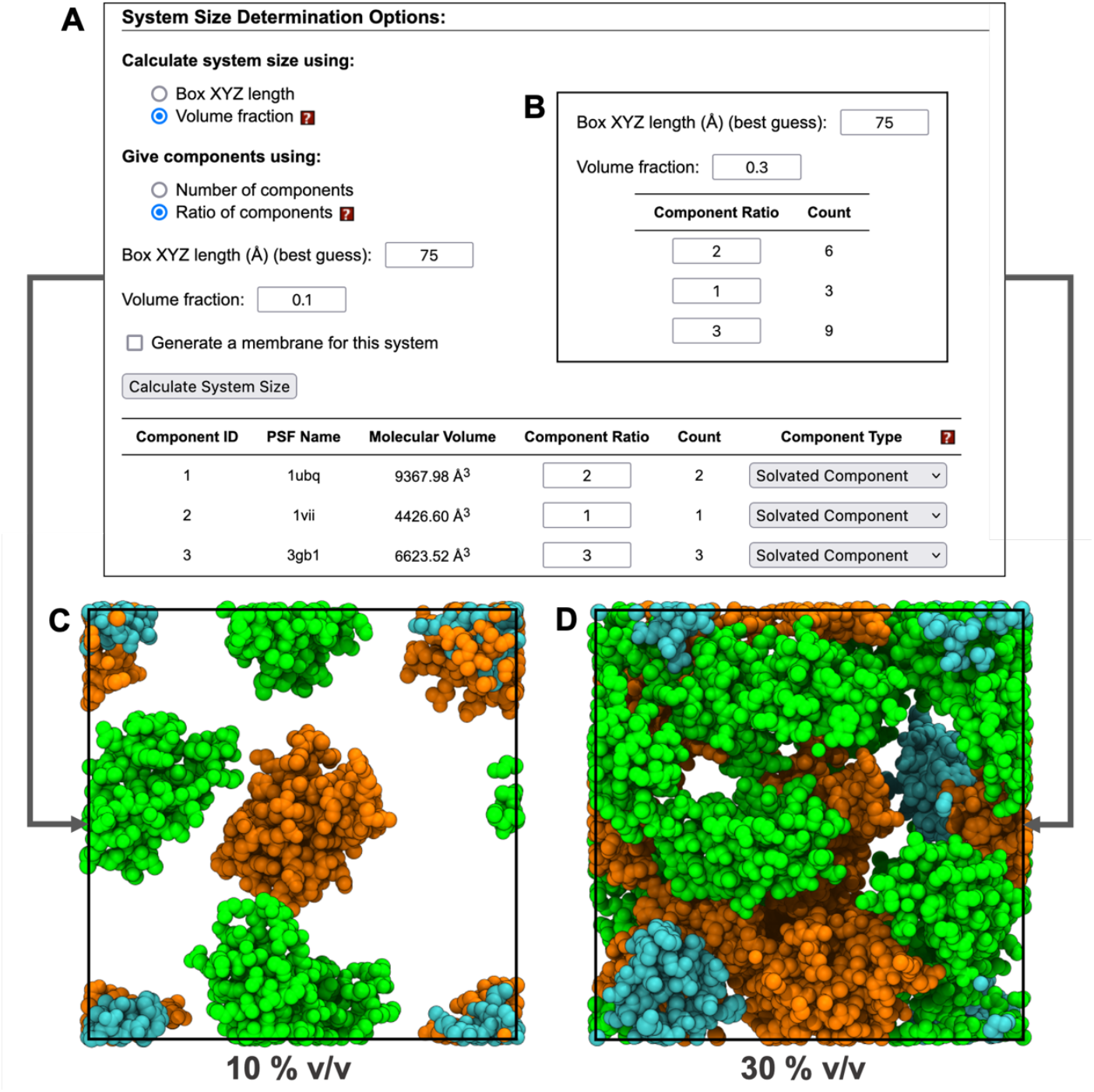
User interface and corresponding packing results with varied volume fraction and component ratios. 10 % v/v is used in (A) and 30 % v/v in (B). Both parameters result in a cube with a side length of 74.19 Å. The resulting component count is updated after clicking “Calculate System Size”. Packing results of (A) and (B) are shown in (C) and (D), respectively. Molecules may cross the system boundaries only if collisions with periodic image atoms are avoided. In (C) and (D), all atoms are wrapped to the primary cell to illustrate the use of available space. Ubiquitin (PDB: 1UBQ), villin (PDB: 1VII), and protein G (PDB: 3GB1) are colored in orange, green, and cyan, respectively.

After the system dimensions and component counts are determined, the user can specify positional constraints (**Figure 4**) whose options differ slightly between component types. For solvated components, the options are: (1) “none”, which accepts any collision-free position and orientation, (2) “planar (Z) restraint”, which fixes the center of mass (COM) of the component to a given Z position, but allows translation along X and Y, and allows any rotation about the component’s COM, and (3) “fixed XYZ”, in which the component’s coordinates are translated so that its COM lies at a given coordinate, and no further movement is allowed. For membrane-embedded components, the options are: (1) “none”, in which the component’s Z positions remain unchanged from their uploaded values, but the component may translate in the X and Y directions and rotate about the Z axis, (2) “planar (Z) restraint”, which is like “none”, but the Z position may be changed a bit from its uploaded value, and (3) “fixed XY”, in which the Z position remains at its initial value, but the user can specify a fixed X and Y coordinate for the component’s COM. Finally, for XY periodic components, the user can specify a COM-Z position, since the X and Y length of periodic components already define the system’s length in X and Y. If no options are selected, the largest component is fixed at the system center. In all cases, the default constraint type is “none”.

**Figure 4.**
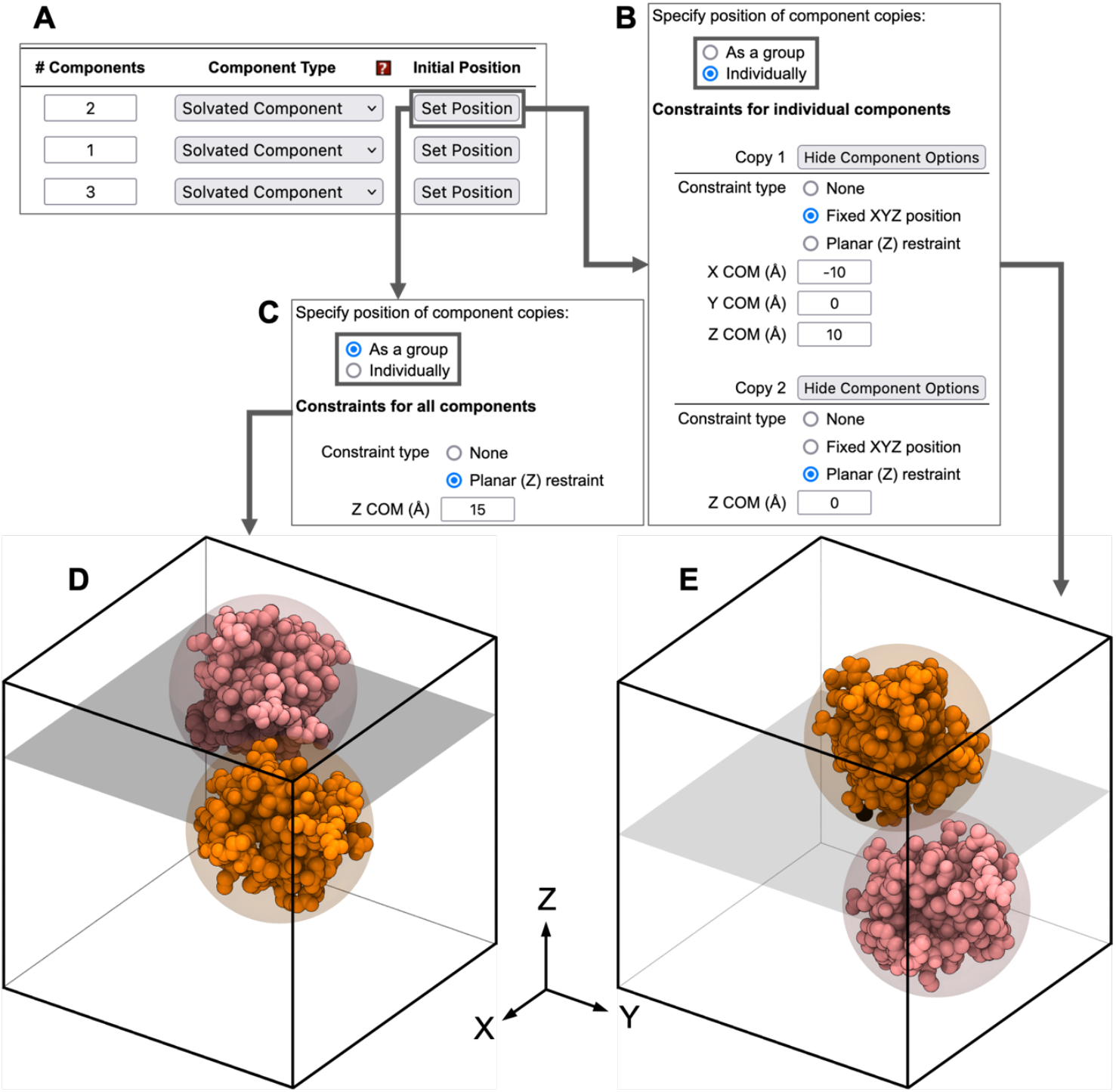
User interface of positioning options for solvated components. Any component can have its positioning constraints modified by clicking “Set Position”, as shown in (A). Exact positions can be specified for (B) individual components’ COM, or (C) identical planar restraints for all copies. Results of applying the settings in (C) and (B) are shown in (D) and (E), respectively. A translucent plane shows the location of each planar restraint, which is applied to both component copies in (D), and only the pink copy in (E). Corresponding coarse-grained (CG) particles are shown as translucent spheres.

The non-solvent components are first packed using an uncharged coarse-grained (CG) sphere model that uses 1 to 3 large atoms per copy of each component (**Figure 5**). Positions are initialized according to the aforementioned positioning constraints chosen by the user. The CG model is then simulated in 5 iterations with Langevin dynamics for 100 ps using a 2 fs timestep at 500 K with a switching function applied to nonbonded forces at 10 Å plus the largest sphere’s radius of gyration. Each iteration of dynamics uses a system size decreasing from 150% of the user’s target to 100%. After dynamics, the CG models are replaced with all-atom models and collisions are minimized with up to 7 iterations of the greedy conformation search (see Supporting Information **Algorithm 1)**.

**Figure 5.**
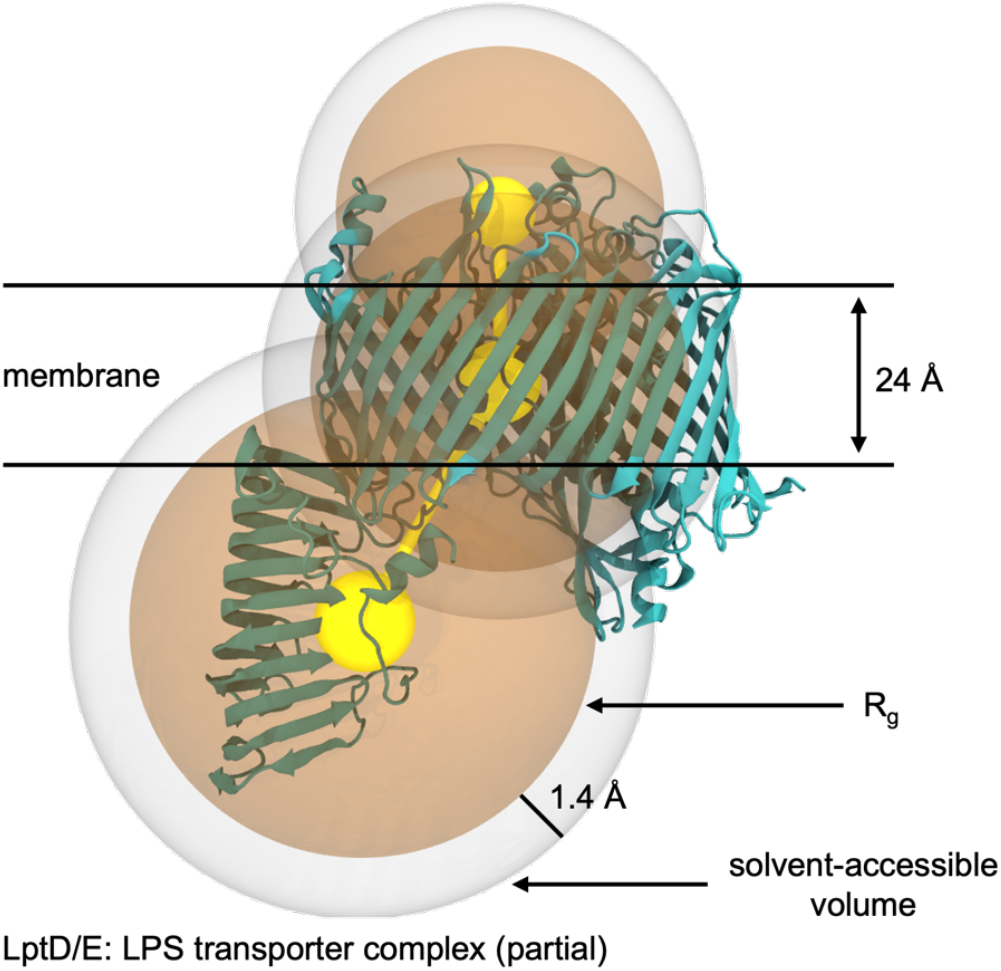
Coarse-grained representations of molecular components. For solvated components, a single particle represents the whole component; for membrane-embedded components, 1-3 particles are used, corresponding to the above-membrane, intra-membrane, and below-membrane regions of the component in a 24 Å thick implicit membrane region centered at Z=0. The van der Waals radius of each particle is set to the radius of gyration (R_g_) of the atoms contained in the corresponding above/intra/below membrane region or omitted if there are no atoms in that region. The solvent accessible volume around a particle corresponds to the 1.4 Å thick shell around that particle’s volume.

If packing fails to find a collision-free configuration, MCA stops and emits an error. Otherwise, segments are renamed using the map from STEP 1. Individual copies of a component are distinguished by appending a copy ID to the segment ID, so that the first protein segment’s copy becomes PAA1, etc.

#### STEP 3 – Build membrane (membrane systems only)

If the system contains membrane-embedded components or the user chooses to generate a membrane, the membrane lipid composition is determined using *Membrane Builder*^38,57,58^ with the system dimensions determined in STEP 2. The lipid packing and replacement procedures are the same as in *Membrane Builder*, except that the system dimensions during lipid packing are decreased in 4 iterations from 150% to 120% of target membrane width, and with 21 iterations from 120% to 100%.

#### STEP 4 – Build solution

The water and ion building procedures in this study follow the same protocols as *Solution Builder*^27,28,43^, with the additional ability to utilize user-uploaded ions. Users can build any combination of ions supported by the CHARMM36 FF, containing up to 7 atoms, in addition to uploading custom ions. Solvent options allow users to specify either the concentration of any solvent, including water, or a solvent density and ratio of solvent volumes. The solvent options page estimates the number of each generated solvent and ion with the user’s settings, although the exact number may differ as described below in STEP 5.

To maintain the approximate solvent ratios, user-uploaded solvent molecules are initially packed into a small box whose size is chosen based on heuristics. The packing procedure follows the same protocols described in STEP 2. After packing, the box is replicated to reach the target box size, and any molecules outside the system boundaries are deleted.

#### STEP 5 – Assemble system

The solvent and non-solvent components are combined by superimposing the solvents with the non-solvents and deleting any solvent molecules that collide with non-solvent molecules within 2.8 Å. If the system contains a membrane or XY periodic component, solvents located in the component’s interior (as defined by the user in STEP 2) are also deleted. If a solvent is present in higher concentration than the user’s target, extras are deleted randomly as needed. However, extra solvent molecules are not created to fill space when system assembly results in fewer than the target number of molecules. Finally, multiple MEMB segments (if any) are joined to a single segment, so that appropriate restraints can be generated in STEP 6.

#### STEP 6 – Generate simulation inputs

MCA uses *FF-Converter* to generate simulation inputs for various MD engines^27,28^. For systems containing periodic nanomaterials or polymers, the generated NVT equilibration inputs use a multi-step scheme where protein restraints are progressively released. Inputs generated for NPT production use anisotropic pressure coupling, and NPAT production inputs use semi-isotropic pressure coupling with pressure applied only along the Z dimension.

### 2. Protein-protein and protein-membrane contacts

The effects of protein crowding near surfaces are important in biology and industry. The prevalence of membrane surfaces in cells suggests that even cytosolic proteins must interact with membranes, especially in organelles with high surface area such as the endoplasmic reticulum and mitochondria. In laboratory settings, solvated proteins can bind to their container surfaces and leave residues that require strong acids to remove. Nawrocki *et al*. recently reported the contacts between crowded proteins and a membrane of cholesterol (CHL1), 1-palmitoyl-2-oleoyl-phosphocholine (POPC), and palmitoyl-sphingomyelin (PSM) via atomistic MD simulation^59^. Protein crowding is difficult to model in atomic details due to the high possibilities for collisions between atoms that cannot be resolved by energy minimization. To study how different surfaces affect protein contact behaviors in crowded environments and to demonstrate the versatility of MCA’s modeling capabilities, we reproduced each of the models in Nawrocki *et al*. and modeled 3 additional membrane types with varying protein concentration.

For the models containing proteins and a membrane-like component (see Methods), we measured the contact probabilities (p) between all pairs of components and aggregated them by component type (**Figure 6**). Our results show that protein contact preferences depend greatly on the membrane environment and protein volume fraction. At 5% v/v, proteins are very likely (p > 0.5) to contact HAP and EO_40_EE_37_ but unlikely (p < 0.1) to contact CHL1/POPC/PSM membranes. Only ubiquitin and villin show high contact probabilities near the axolemma membrane at 5% v/v. For all membrane types, increasing protein v/v has the effect of increasing the relative proportion of contacts between proteins and other proteins, though contacts between protein and membrane remain frequent (p > 0.14) for all protein types and membranes except CHL1/POPC/PSM.

**Figure 6.**
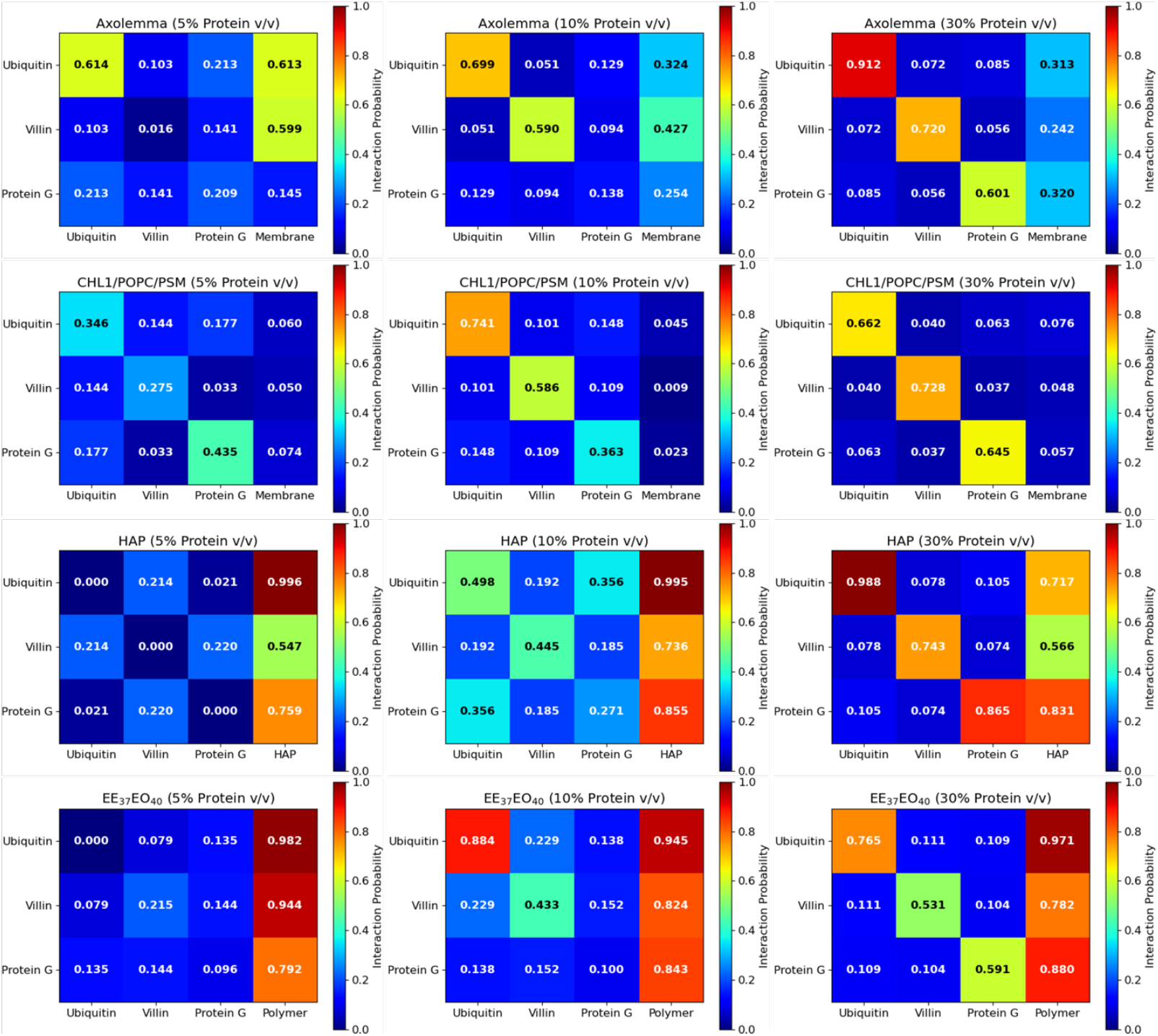
Protein contact probability for all systems containing membrane-like components. A contact is defined when Cα-Cα distance ≤ 7 Å. Note that for calculation of % v/v in CHL1/POPC/PSM systems only, protein volume includes the solvent accessible surface area (SASA).

Proteins uniformly prefer contacts with the same type of protein than with other protein types, with two notable exceptions. First, at 5% protein v/v, villin prefers contacts with axolemma membrane over contacts with other copies of villin. Second, no obvious contact patterns between proteins are observed near HAP for protein v/v < 30% because contacts with HAP are strongly preferred.

Visualization of the HAP and EO_40_EE_37_ simulation trajectories suggests different reasons for their frequent protein contacts. The duration of contact events between proteins and HAP is higher than that in other membranes, with several contact events lasting longer than 100 ns (**Figure S4**). In contrast, the hydrophilic EO domain of EO_40_EE_37_ dissolved so much into water throughout simulation that there was almost no region of the solution where EO was absent (**Figure S5**) as shown by the width of EO_40_EE_37_’s distribution in **Figure S1**. This behavior suggests that EO_40_EE_37_ should be equilibrated separately with a thick water layer before being combined with proteins via MCA.

### 3. POPC diffusion near mica-supported lipid bilayers

Planar-supported lipid bilayers (SLBs), artificial membranes consisting of a lipid bilayer deposited on a solid support, have become an essential tool for investigating the properties and functions of biological membranes. Compared to other membrane models, such as giant unilamellar vesicle or cell membranes, SLBs offer numerous advantages, including their simplicity, reproducibility, and versatility in analytical techniques, such as fluorescence microscopy, surface plasmon resonance, and atomic force microscopy^60^. Therefore, SLBs have been widely utilized in the fields of biophysics, biochemistry, and cell biology to study a variety of membrane-related processes, such as membrane protein function, protein-lipid interactions, and membrane fusion. Therefore, understanding the lateral motion of lipids in SLBs is crucial in elucidating the properties and functions of biological systems. The lateral mobility of lipids is a key determinant of membrane structure and function, and is influenced by various factors such as temperature, lipid composition, and the presence of membrane proteins. Thus, accurate measurements of lipid diffusion coefficients are necessary to fully understand membrane behavior and corresponding biological events.

In this study, we investigated the effect of the solid support on lipid diffusion in SLBs using model systems (**Figure 1E**). We prepared three systems with a pure POPC bilayer separated by 1, 2, and 3 nm from a mica support, and analyzed the diffusion coefficient of POPC lipids in the lower (D_B-1nm_, D_B-2nm_, and D_B-3nm_) and upper (D_T-1nm_, D_T-2nm_, and D_T-3nm_) leaflets using the Diffusion Coefficient Tool^61^ plugin in VMD^34^. Our MD simulation results (**Figure 7**) show that D_B-1nm_, D_B-2nm_, and D_B-3nm_ are 4.8 μm^2^/s, 6.6 μm^2^/s, and 6.4 μm^2^/s, respectively (all errors are less than 0.01 μm^2^/s). Experimental values of the POPC diffusion coefficient in a bilayer are ∼7 μm^2^/s^62^, implying that a bilayer on support behaves like free-standing lipids when the water thickness is more than 2 nm. In contrast, when the thickness of the water layer is less than 2 nm, there are strong interactions between the support and the lipids, which reduces the diffusion coefficient of the lipid in the lower leaflet of the SLB. This finding is consistent with experimental measurements, which have shown that the thickness of the water layer is in the range of 1∼2 nm^62,63^. In addition to the lower leaflet, D_T-1nm_, D_T-2nm_, and D_T-3nm_ show 5.9 μm^2^/s, 6.7 μm^2^/s, and 6.5 μm^2^/s, respectively (all errors are less than 0.01 μm^2^/s). These results suggest that the interaction between the mica support and lower leaflet affects the diffusion of lipids in the upper leaflet as well. The decrease in D_T-1nm_ is likely due to the coupling of the two leaflets via lipid-lipid interactions across the lipid bilayer. This finding highlights the importance of considering the behavior of both the upper and lower leaflets when studying the properties and functions of biological membranes in contact with solid supports.

**Figure 7.**
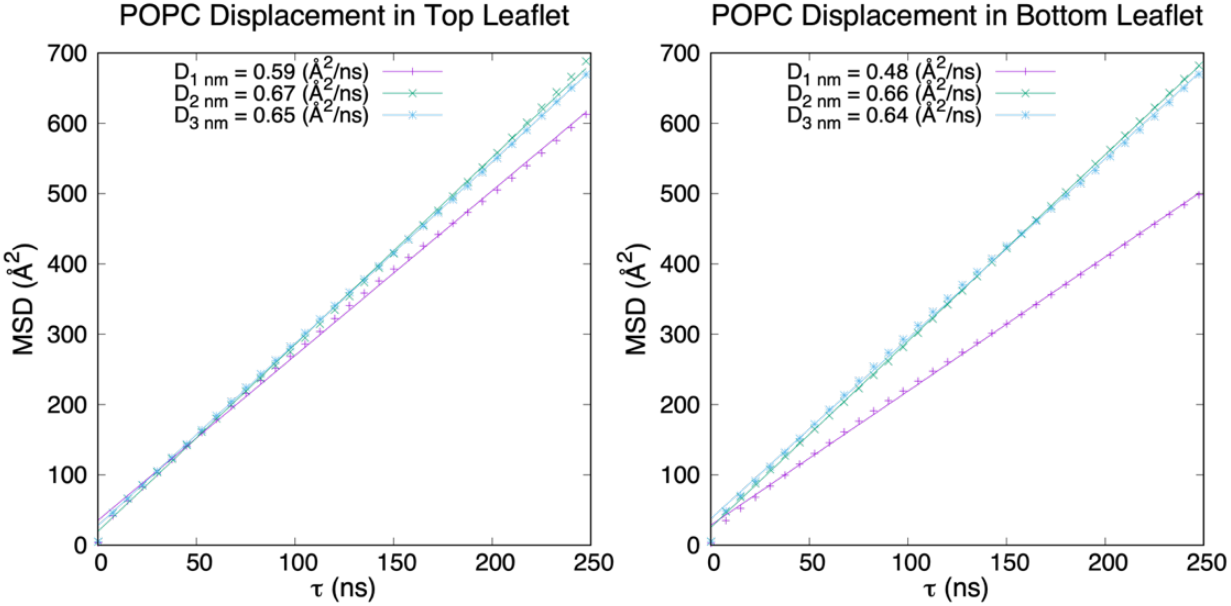
POPC diffusion near mica-supported lipid bilayers. Upper and lower leaflets are distal and proximal to mica, respectively. Mean square displacement (MSD) of POPC phosphorus is plotted against lag time, with diffusion coefficient (D) shown as a fitted line. All lag times between 1 to 250 ns were used to calculate D.

### 4. Diffusion of CO_2_ through polymer membranes

Polyethylene terephthalate (PET) is a cheap, recyclable plastic used in containers, clothing, and many other products with the resin ID code 1. Modern industrial production and recycling of PET causes significant non-renewable energy usage (NREU) and release of greenhouse gases (GHGs)^64^, which has led to an investigation into potential replacement plastics that could be constructed from renewable sources. One such candidate is poly(ethylene 2,5-furandicarboxylate) (PEF), which is amenable to production from biomass and which could substantially reduce NREU and GHG production^65^. In experimental investigations of the permeability of plastics to CO_2_ and other gases, PEF has shown uniform improvement over PET in its CO_2_ retention ability^66^. To investigate possible mechanisms for CO_2_ diffusion through PET and PEF, we simulated 100 Å thick polymer sheets of each plastic with 1 atm initial pressure in pure CO_2_ (**Figure 1F**).

Our results show that CO_2_ penetration in PEF_95_ is comparable to that in PET_95_ at each measured simulation time point. In our simulations, CO_2_ molecules made rapid jumps between defects located at varying depths in the polymer structures, followed by long periods where the molecules remained in the same approximate region. The exact location of these defects varies greatly between replicas, as indicated by the error bars in **Figure 8**, but the general trend for both polymers is an increase in overall CO_2_ diffusion over time. Notably, between 666–1333 ns, the CO_2_ density in near the polymer center (0–12 Å) was higher for PET_95_ than for PEF_95_, and the CO_2_ density near the polymer periphery (15–37 Å) was lower for PET_95_ than for PEF_95_ in the same time range, indicating that PEF_95_ is overall more resistant to CO_2_ diffusion.

**Figure 8.**
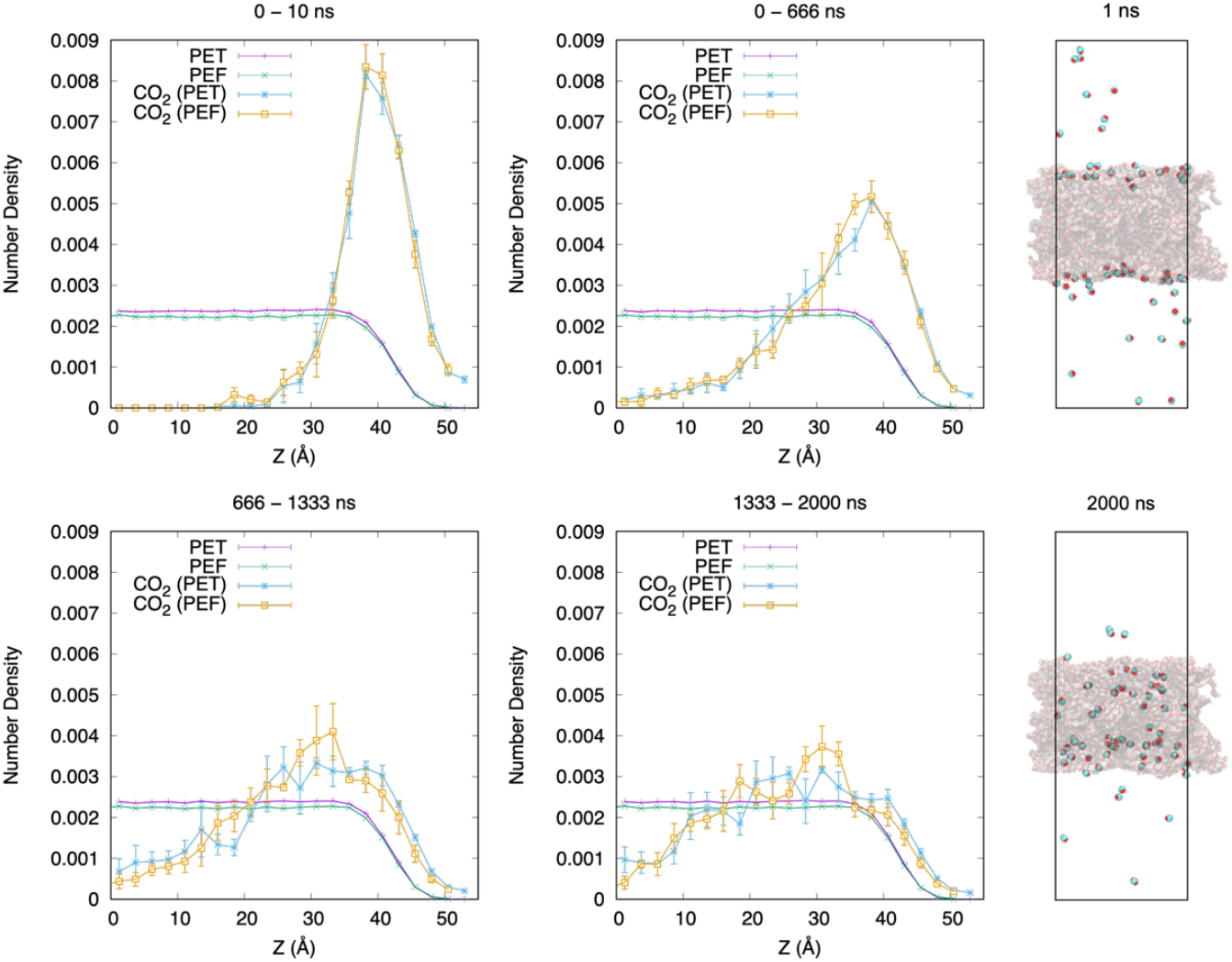
Symmetrized Z density profiles of CO_2_ in PET_95_ and PEF_95_. Left: mean ± standard error of replicas of the same plastic are plotted with a bin size of 2.46 Å. PEF_95_ and CO_2_ in PEF_95_ values are shifted left by 2.46 Å to correct for changes in plastic thickness that occurred during equilibration in vacuum. Bin heights are normalized to sum to 1. Right: simulation snapshots of PET_95_ and CO_2_ taken after 1 ns and 2000 ns of NVT simulation.

## Conclusions

This work presents an automated and versatile procedure for building simulation-ready, atomistic models containing heterogeneous molecular components via *Multicomponent Assembler* (MCA) in CHARMM-GUI. Automation is achieved by assigning component types that determine the packing strategy and using a greedy packing algorithm whose initial configuration is generated from coarse-grained simulation. The provided positioning options allow constraining components to a given position or orientation while allowing them to move within those constraints during packing.

Although MCA requires that input structures are provided in CHARMM format, other CHARMM-GUI tools—especially *PDB Reader & Manipulator*^55^, *Ligand Reader & Modeler*^21^, *Glycan Reader & Modeler*^56^, *Nanomaterial Modeler*^44^, and *Polymer Builder*^45^—can be used to prepare these initial structures. Furthermore, its integration with *FF-Converter*^27,28^ allows MCA to provide simulation inputs using both CHARMM and AMBER FFs and for many other simulation programs, including NAMD^33^, GROMACS^31,32^, Amber^30^, and OpenMM^35^. MCA’s diverse modeling capabilities are demonstrated by building crowded solute-membrane interfaces, nano-bio interfaces, multi-membrane systems, and non-water solvents. Our modeling of polymer-bio and nano-bio interfaces are made possible by the CHARMM36(m)^46^ and INTERFACE FFs^49–54^ and greatly facilitated by *Polymer Builder*^45^ and *Nanomaterial Modeler*^44^.

To the best of our knowledge, this work is the first to solve a PBC-aware packing problem for large, densely packed molecules. In combination with other CHARMM-GUI modules, MCA can generate complex molecular systems by combining many components. We hope that MCA can facilitate innovative studies of complex interactions between small (organic and inorganic) molecules, biomacromolecules, polymers, and nanomaterials.

## Methods

### Test System Preparation

To test MCA’s ability to generate complex multicomponent systems reliably, we have built and simulated the systems shown in **Figure 1**. Each system’s configuration is summarized in **Table S1** and the preparation is described below. Video demos of MCA usage for most of the below molecular configurations are available at https://charmm-gui.org/demo. The FFs necessary for nanomaterial and polymer modeling^49–54^ are automatically included in the systems prepared by CHARMM-GUI. Note that protein-water interaction scaling previously used to reduce protein-protein interactions^67^ was not used in this study.

#### 1. Solution systems with three proteins (5, 10, 30% v/v)

We built three systems described by Nawrocki *et al*.^59^, each containing ubiquitin (PDB ID: 1UBQ), villin (PDB ID: 1VII), and streptococcal G protein (PDB ID: 3GB1) in volume fractions (v/v) of 5%, 10%, and 30%, solvated by TIP3P water and 150 mM KCl (**Table S1**). Each system was simulated with OpenMM using the NVT (constant particle number, volume, and temperature) equilibration and NPT (constant particle number, pressure, and temperature) production protocols provided by CHARMM-GUI *Solution Builder*^27,28,43^ at 310.15 K with hydrogen mass repartitioning (HMR)^68^ for 1 μs.

##### 2-1. Membrane systems with three proteins (5, 10, 30% v/v)

We built three systems containing the same proteins in Section 1, combined with a membrane containing an equal ratio of cholesterol (CHL1), 1-palmitoyl-2-oleoyl-phosphatidylcholine (POPC), and palmitoylsphingomyelin (PSM); see **Table S1** for system information. Membranes were built *de novo* in MCA using the *Membrane Builder* protocol in STEP 3. Systems were simulated with OpenMM using the multi-step NVT/NPT equilibration and NPT production protocols provided by *Membrane Builder*^38,39^ at 310.15 K with HMR for 1 μs. As the CHL1/POPC/PSM membrane system was not pre-equilibrated, its size changed significantly throughout the first 100 ns of simulation. In each system, the X and Y axes shrank while the Z axis grew (**Figure S3**).

##### 2-2. Pre-equilibrated axolemma membrane systems with three proteins (5, 10, 30 % v/v)

An axolemma membrane model that was simulated by Lee *et al*.^39^ was prepared by manually removing water and ions from the last simulation frame. Its PSF/CRD files were then uploaded with those of 1UBQ, 1VII, and 3GB1. We used the solvated component type for the three proteins and the periodic type for the axolemma membrane. Dimensions for the axolemma were taken from the previous simulation (for X and Y) and the membrane’s Z dimension was estimated as ∼50 Å. **Table S1** summarizes the final system information. To speed up packing for the 5% and 10% v/v cases, protein components were excluded above and below 10 Å from the membrane region. Systems were simulated with OpenMM using the multi-step NVT/NPT equilibration and NPT production protocols provided by *Membrane Builder*^38,39^ at 310.15 K with HMR for 1 μs.

#### 3. HAP with three proteins (5, 10, 30% v/v)

We first constructed a hydroxyapatite (HAP) slab with a dimension of 103.6 Å x 114.2 Å x 42 Å at pH 10 using *Nanomaterial Modeler*^44^. The HAP slab was then uploaded with the same proteins described in Section 1. The solvated component type was used for the three proteins, and the periodic type was used for HAP. To achieve protein volume fractions of 5/10/30, the number of protein copies (2, 4, and 17 each) and system Z dimension were varied (**Table S1**). To speed up packing for all cases, protein components above and below 5 Å from the HAP region were excluded. After system assembly, 5000 steps of steepest descent (SD) minimization was followed by 5000 steps of adopted basis Newton-Raphson (ABNR) minimization using CHARMM. The systems were then simulated without HMR at 303.15 K for 1 μs using OpenMM.

##### 4-1. Polymer EO_40_EE_37_ with three proteins (5, 10, 30% v/v)

We first constructed a polyethylene oxide-poly(ethylethylene) polymer slab (EO_40_EE_37_) in solution using *Polymer Builder*, as described by Choi *et al*.^45^, with a thickness of 120 Å and a width of 107.9 Å. The slab was then uploaded with the same proteins described in Section 1. The solvated component type and periodic type were used for the three proteins and the EO_40_EE_37_, respectively. To achieve protein volume fractions of 5/10/30, the number of protein copies (2, 4, and 13 each) and system Z dimension were varied (**Table S1**). To speed up packing for the 5% and 10% v/v cases, protein components were excluded above and below 10 Å from the polymer region. After system assembly, we performed 5000 steps of SD minimization followed by 5000 steps of ABNR minimization using CHARMM. Systems were then simulated without HMR at 303.15 K for 1 μs.

##### 4-2. Diffusion of CO_2_ through polymer membranes

We used *Polymer Builder* to construct slabs containing polyethylene terephthalate (PET_95_) and polyethylene 2,5-furandicarboxylate (PEF_95_) with a monomer length of 95 each in vacuum with a thickness of 100 Å and width near 100 Å. This resulted in 40 molecules of PET_95_ with a width of 108.4 Å and 46 molecules of PEF_95_ with a width of 108.4 Å. We ran equilibration and 500 ns of production for each slab separately in vacuum using the default inputs provided by *Polymer Builder* at 298.15 K without HMR. We obtained a CO_2_ structure from the CHARMM36 FF by *Ligand Reader & Modeler*. To combine each polymer with CO_2_, we used the solvent component type for CO_2_ and periodic type for the PET_95_ or PEF_95_. Three replicas of each polymer + CO_2_ system were constructed (**Table S1**). Instead of a water solvent, we used the CO_2_ to construct a gaseous solvent with pure CO_2_ present at 1.98 g/L density. This resulted in 64 copies of CO_2_ with PET_95_ and 64 copies of CO_2_ with PEF_95_. After system assembly, we performed 5000 steps of SD minimization followed by 5000 steps of ABNR minimization using CHARMM. Systems were then simulated without HMR at 298.15 K for 2 μs using OpenMM.

#### 5. Multi-layer system (mica + POPC membrane)

To demonstrate the capability of MCA to build multi-layer models, we obtained a mica model of 103.8 Å x 108.2 Å x 76.8 Å from *Nanomaterial Modeler* and uploaded it to MCA using the periodic component type and selecting the option to generate a new membrane. To analyze the effect of mica on the membrane, we generated three such models by varying the Z position of the uploaded mica’s COM, Z_1_ = 48.33 Å, Z_2_ = 58.33 Å, and Z_3_ = 68.33 Å, corresponding to 10 Å, 20 Å, and 30 Å initial separation between membrane head group atoms (initialized near Z = 19 Å) and mica. A pure POPC bilayer containing 165 lipids in each leaflet was constructed with the membrane centered at Z = 0. The systems were solvated with 0.15 M KCl followed by 100 steps of SD minimization in CHARMM and 5000 steps of minimization in OpenMM. **Table S1** summarizes the final system information. The mica + POPC systems were equilibrated in a multistage procedure starting with 250 ps NVT simulation at 298.15 K with positional and dihedral restraints starting at 1000 kJ/mol/nm^2^ and 1000 kJ/mol/rad^2^ decreasing halfway to 400 kJ/mol/nm^2^ and 400 kJ/mol/rad^2^. We then used an NPT ensemble at 298.15 K for 3.625 ns with restraints decreasing from 400 kJ/mol/nm2 and 200 kJ/mol/rad2 to 0 and pressure coupling applied in all dimensions at 1 atm, as shown in **Figure S2**. Finally, all systems were run for 1 μs under NVT ensemble at 298.15 K without HMR.

## Supporting information

MCA-SI

## Acknowledgements

This work has been supported by NIH GM138472.

## Author Contributions

N.K., Y.K.C., J.L., and W.I. designed the research. N.K performed the research, wrote the software, and drafted the paper with input from the co-authors. Y.K.C. wrote the text for POPC diffusion near mica-supported lipid bilayers. J.L. helped with FF Converter integration. Y.K.C. and J.L. recommended appropriate default settings for simulation inputs. Y.K.C., J.L., and W.I. helped with interpretation of results and substantially revised the manuscript.

