## Supplementary material for "CHARMM-GUI Multicomponent Assembler for Modeling and Simulation of Complex Multicomponent Systems": MCA-SI

**Algorithm 1.** Packing optimization. A greedy search of translation and rotation space for a given component (comp). cmax is the maximum allowed number of collisions. dcut is the distance around any atom in comp to check for collisions. $\delta\theta$ is the rotation increment. $\theta_0$ is the initial rotation angle along each axis. $\theta_{max}$ is the maximum rotation angle along each axis. Collision-detection ignores CG particles if any have not yet been replaced.

```
GreedyTransRotSearch(comp, cmax, dcut, δxyz, δθ, θ0, θmax):
    let best = (Infinity)
    if component type is membrane
        for all (tx, ty) in 2-fold Cartesian product of {-δxyz, 0, δxyz}
            translate comp by tx, ty
            if DoRotation(comp, cmax, dcut, δθ, θ0 , θmax, tx, ty, 0, best)
                return
            endif
            undo translation
        endfor
    else
        for all (tx, ty, tz) in 3-fold Cartesian product of {-δxyz, 0, δxyz}
            translate comp by tx, ty, tz
            if DoRotation(comp, cmax, dcut, δθ, θ0, θmax, tx, ty, tz, best)
                return
            endif
            undo translation
        endfor
    endif
    // if this line reached, no trans/rot is ≤ cmax; restore best trans/rot
    translate comp by best[1], best[2], best[3]
    rotate comp by best[4], best[5], best[6]
endfunc

DoRotation(comp, cmax, dcut, δθ, θ0, θmax, tx, ty, tz, best):
    let angles = {}
    for i=0 to floor((θmax - θ0) / δθ)
        append (θ0 + i*δθ) to angles
    endfor
    if component type is membrane
        angles = {(0, 0, θz) forall θz in angles}
    else
        angles = 3-fold Cartesian product of angles
    endif
    forall (θx, θy, θz) in angles
        rotate comp to (θx, θy, θz) along (x, y, z) axes
        let collisions = number of atoms of comp within dcut of
                  atoms not in cmax, including image atoms
        if component type is not membrane
            collisions += number of atoms of comp within dcut of
                   membrane exclusion region
        endif
        if collisions < best[0]
            best = (collisions, tx, ty, tz, θx, θy, θz)
        endif
        if collisions ≤ cmax then return true
    endfor
    undo rotation
    return false
endfunc
```

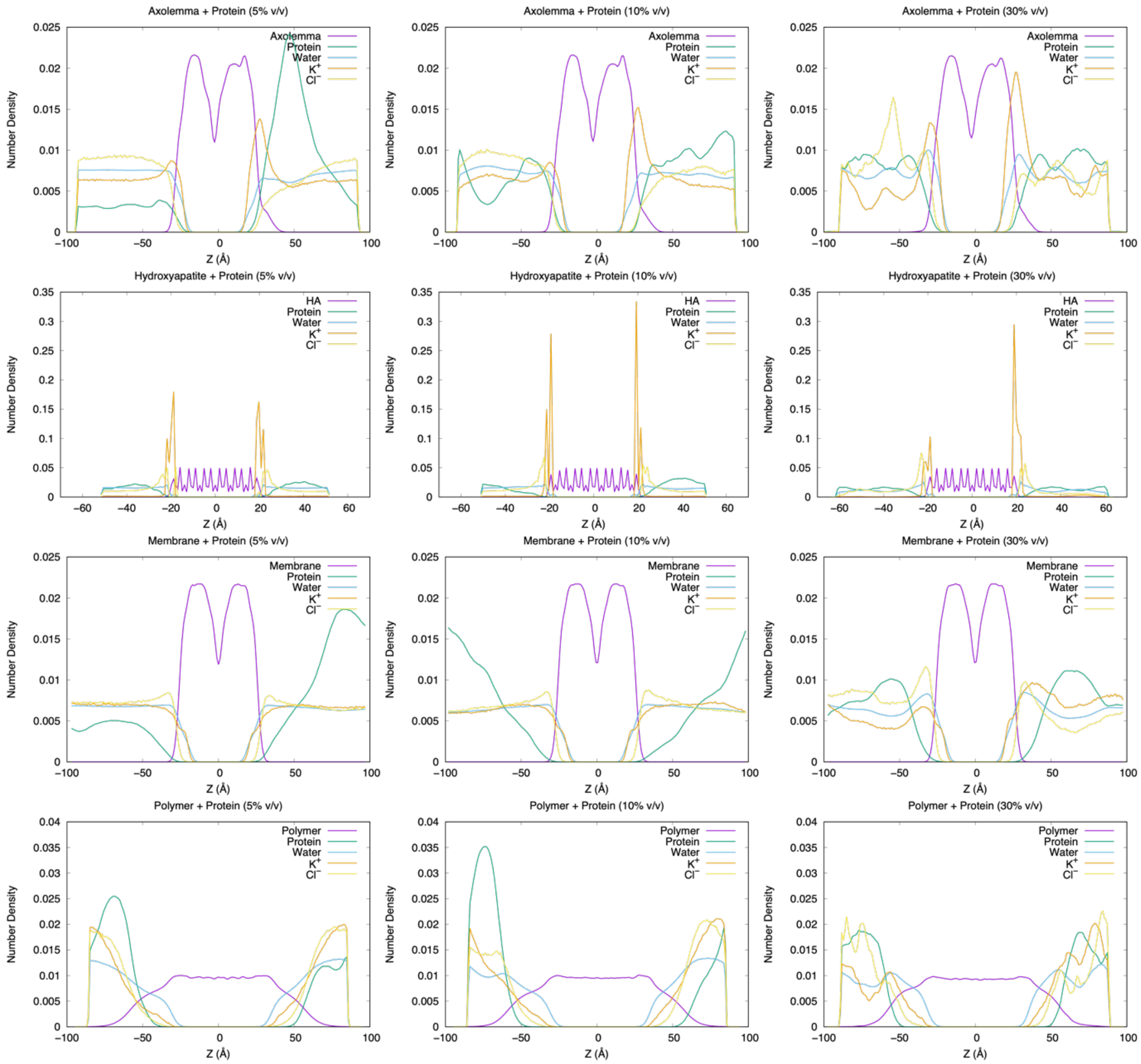


**Figure S1.** Z density profiles of all systems containing a membrane-like component and proteins. Values are averaged across the whole simulation and plotted with a bin size of 1 Å. Bin heights are normalized to sum to 1.

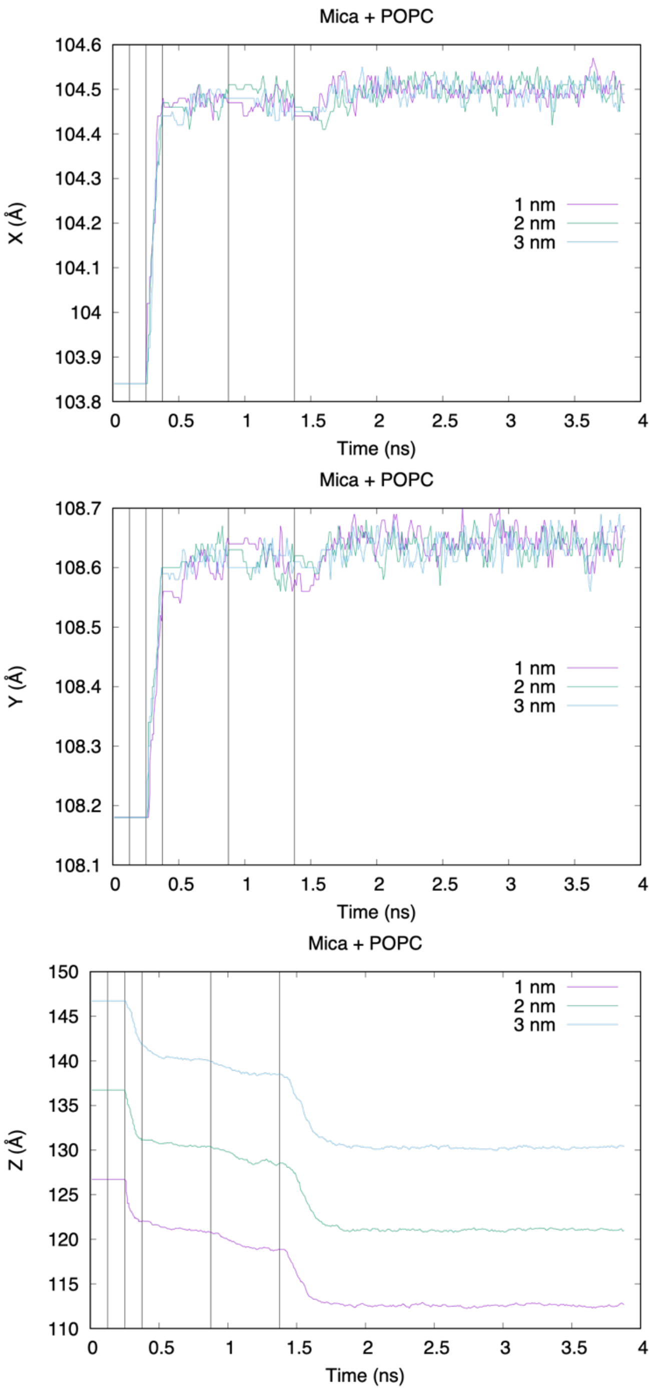


**Figure S2.** System dimensions of mica + POPC systems during equilibration. Vertical bars indicate the simulation times at which the strength of restraints was decreased.

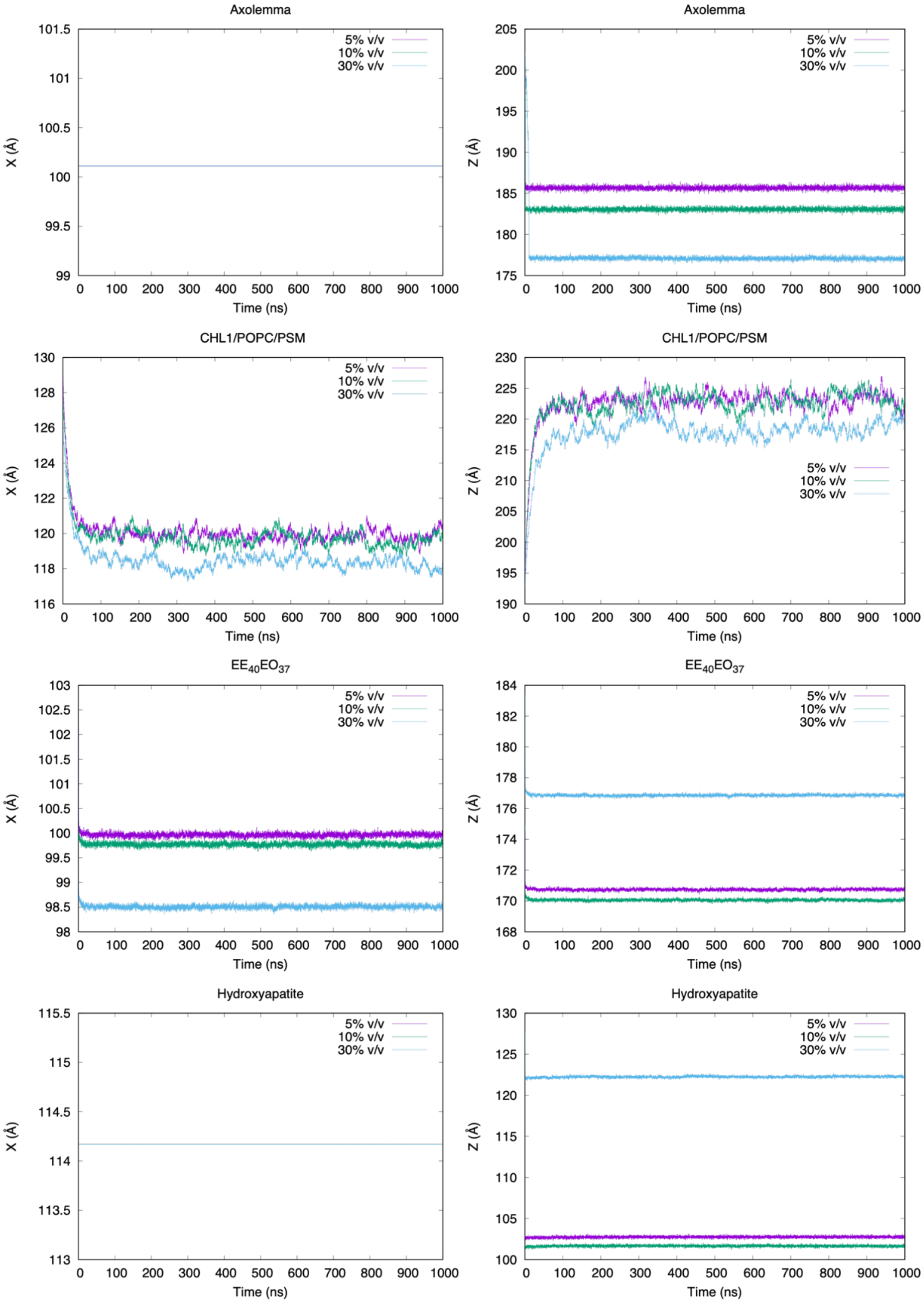


**Figure S3.** System dimensions of all systems containing a membrane-like component and proteins during production runs. Y dimensions are omitted because X = Y.

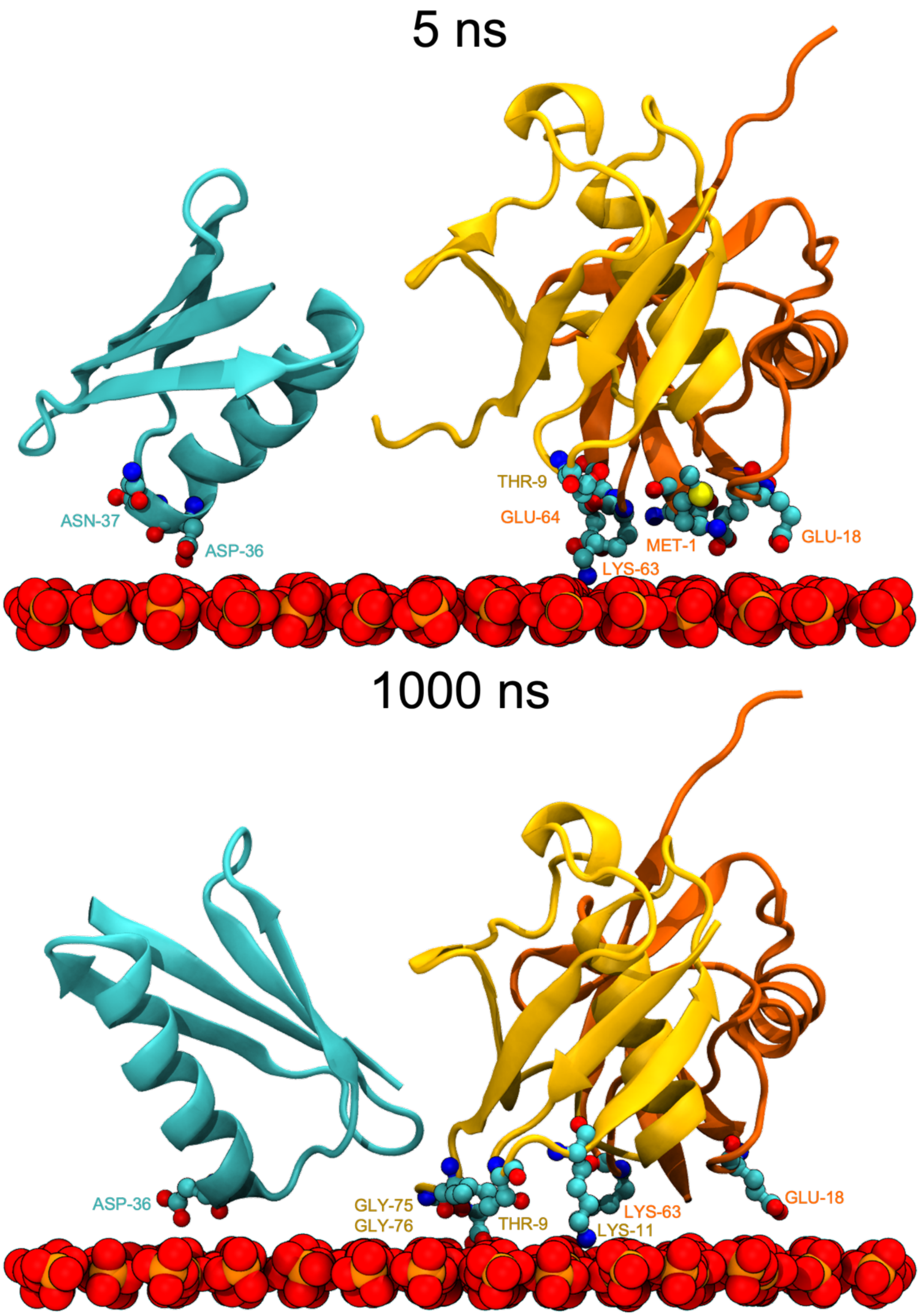


**Figure S4.** Example protein cluster with long-lived contact to HAP. The contact event shown begins at 5 ns and lasts for the rest of the 1 μs simulation. The cluster consists of one protein G (PDB: 3GB1, cyan) and two ubiquitin (PDB: 1UBQ, yellow and orange). Proteins in this system occupied 10% of solvent volume.

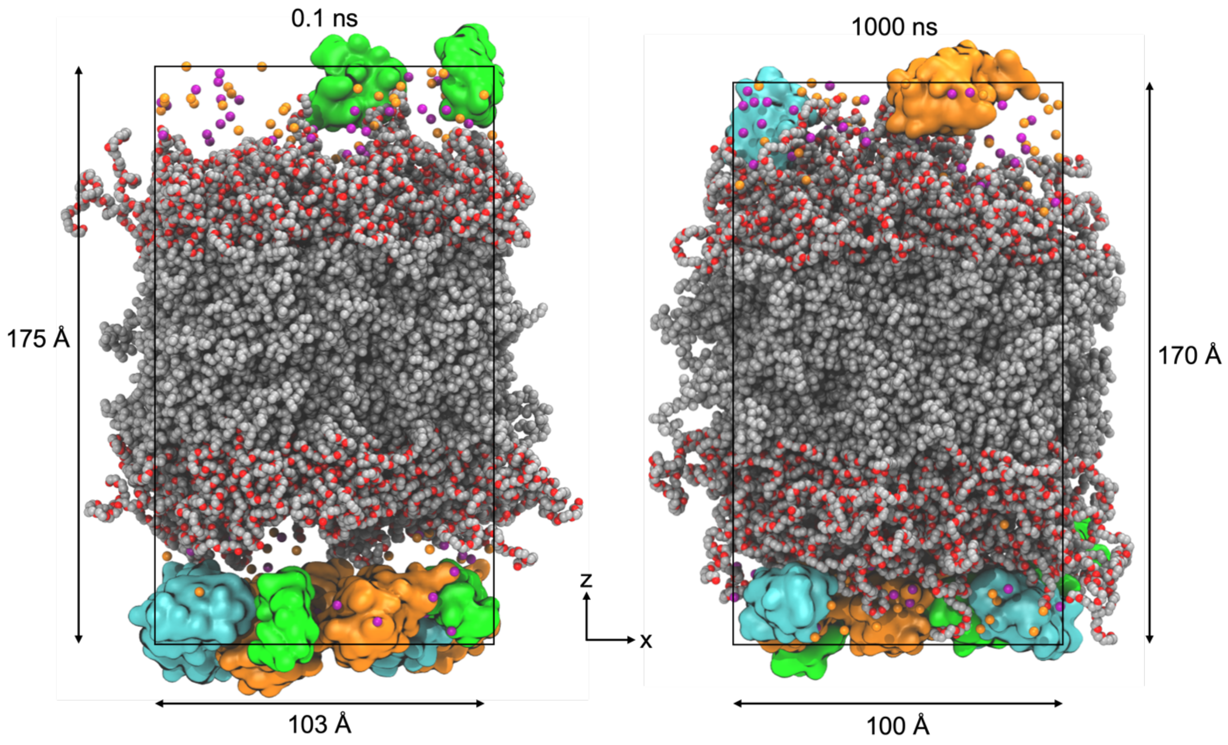


**Figure S5.** Initial and final snapshots of $EO_{40}EE_{37}$ polymer membrane with proteins. The color scheme is the same as in Figure 1D.

**Table S1.** System configurations used in this study.

| System Type | Protein v/v | Component | Copies | Initial System Dimensions | Production Runtime |
|---|---|---|---|---|---|
| Solvated Proteins | 5% | 1UBQ | 5 | (153.32 Å)$^3$ | 1 µs |
| | | 1VII | 5 | | |
| | | 3GB1 | 5 | | |
| | 10% | 1UBQ | 10 | (154.06 Å)$^3$ | 1 µs |
| | | 1VII | 10 | | |
| | | 3GB1 | 10 | | |
| | 30% | 1UBQ | 10 | (106.31 Å)$^3$ | 1 µs |
| | | 1VII | 10 | | |
| | | 3GB1 | 10 | | |
| Proteins + Membrane | 5% | 1UBQ | 4 | (134.21 Å)$^2$ x 193.71 Å | 1 µs |
| | | 1VII | 4 | | |
| | | 3GB1 | 4 | | |
| | | Cholesterol | 110:111 | | |
| | | POPC | 110:111 | | |
| | | PSM | 110:111 | | |
| | 10% | 1UBQ | 8 | (134.18 Å)$^2$ x 195.34 Å | 1 µs |
| | | 1VII | 8 | | |
| | | 3GB1 | 8 | | |
| | | Cholesterol | 110:110 | | |
| | | POPC | 110:110 | | |
| | | PSM | 110:110 | | |
| | 30% | 1UBQ | 23 | (133.74 Å)$^2$ x 194.83 Å | 1 µs |
| | | 1VII | 23 | | |
| | | 3GB1 | 23 | | |
| | | Cholesterol | 108:107 | | |
| | | POPC | 108:107 | | |
| | | PSM | 108:107 | | |
| Proteins + Axolemma | 5% | 1UBQ | 5 | (100.11 Å)$^2$ x 202.99 Å | 1 µs |
| | | 1VII | 5 | | |
| | | 3GB1 | 5 | | |
| | | Axolemma | 1 | | |
| | 10% | 1UBQ | 10 | (100.11 Å)$^2$ x 202.99 Å | 1 µs |
| | | 1VII | 10 | | |
| | | 3GB1 | 10 | | |
| | | Axolemma | 1 | | |
| | 30% | 1UBQ | 10 | (100.11 Å)$^2$ x 206.2 Å | 1 µs |
| | | 1VII | 10 | | |
| | | 3GB1 | 10 | | |
| | | Axolemma | 1 | | |

| System Type | Protein v/v | Component | Copies | Initial System Dimensions | Production Runtime |
| --- | --- | --- | --- | --- | --- |
| Proteins + HAP | 5% | 1UBQ<br>1VII<br>3GB1<br>HAP | 2<br>2<br>2<br>1 | 103.59 x 114.17 x 111.06 Å$^3$ | 1 μs |
| | 10% | 1UBQ<br>1VII<br>3GB1<br>HAP | 4<br>4<br>4<br>1 | 103.59 x 114.17 x 111.06 Å$^3$ | 1 μs |
| | 30% | 1UBQ<br>1VII<br>3GB1<br>HAP | 17<br>17<br>17<br>1 | 103.59 x 114.17 x 111.06 Å$^3$ | 1 μs |
| Proteins + $EO_{40}EE_{37}$ | 5% | 1UBQ<br>1VII<br>3GB1<br>$EO_{40}EE_{37}$ | 2<br>2<br>2<br>1 | (107.88 Å)$^2$ x 189.51 Å | 1 μs |
| | 10% | 1UBQ<br>1VII<br>3GB1<br>$EO_{40}EE_{37}$ | 4<br>4<br>4<br>1 | (107.88 Å)$^2$ x 189.51 Å | 1 μs |
| | 30% | 1UBQ<br>1VII<br>3GB1<br>$EO_{40}EE_{37}$ | 13<br>13<br>13<br>1 | (107.88 Å)$^2$ x 196.04 Å | 1 μs |

| System Type* | $CO_2$ Density | Component | Copies | Initial System Dimensions | Production Runtime |
| --- | --- | --- | --- | --- | --- |
| $PET_{95}$ + $CO_2$ | 1.98 g/L | $PET_{95}$<br>$CO_2$ | 1<br>64 | (108.4)$^2$ x 300.00 Å | 2 μs |
| $PEF_{95}$ + $CO_2$ | 1.98 g/L | $PEF_{95}$<br>$CO_2$ | 1<br>64 | (108.4 Å)$^2$ x 300.00 Å | 2 μs |

*3 replicas of each PET and PEF system were built with the same parameters.
